# Stratification in Microbial Communities with Depth and Redox Status in a Eutrophic Lake Across Two Years

**DOI:** 10.1101/2021.10.15.464574

**Authors:** Robert A. Marick, Benjamin D. Peterson, Katherine D. McMahon

## Abstract

Bacteria have a profound impact on many key biogeochemical cycles in freshwater lake ecosystems; in turn, the composition of bacteria in the lake is contingent on the chemistry of the water. Many parameters that affect bacterial growth in freshwater ecosystems, such as water temperature, nutrient levels, and redox status, exhibit notable inter-annual differences in addition to seasonal changes. However, little is known about the impact of these inter- and intra-annual differences on the freshwater microbiome, especially in anoxic bottom waters. In this study, we paired biogeochemical field data with 16S rRNA gene amplicon sequencing of depth-discrete samples from a dimictic lake across two open-water seasons to observe variation in the microbiome relative to differences in water chemistry between two years. We found differences in the timing anoxia onset and the redox status in the water column across the two years. Changes in redox status led to major shifts in the microbial community composition. While there was little variation between years in the microbial taxonomic composition at the phyla level, there was substantial interannual variation at more resolved taxonomic levels. Some interannual differences can be explained by links between the predicted metabolic potential of those lineages and the different redox conditions between the two years. These results emphasize the need for repeated monitoring to deduce long-term trends in microbial communities in natural ecosystems and the importance of a comprehensive evaluation of environmental conditions contemporary with any microbiome analysis.

**Importance:** The results of this study add to the growing body of evidence that microbial communities in natural systems are temporally dynamic on multiple scales, and even more so at highly resolved taxonomic levels. By correlating our analysis of the microbial community with the redox status of the water column we find that many community differences between the years can be in part explained by these parameters. As collecting 16S rRNA data over many years is critical to understanding long term trends in microbial ecology, our study suggests that corresponding water chemistry data could be a powerful tool to help explain microbiome trends.

## Introduction

Bacterial communities have a significant impact on the ecosystems in which they reside due to the domain’s diverse range of metabolisms that interact with virtually all biogeochemical cycles (Falkowski, Fenchel, and Delong 2008). A key goal of microbial ecology is to better understand and predict these bacterial communities’ biodiversity and how this diversity is governed by abiotic environmental factors. In aquatic environments, many different chemical and physical properties of the water are known to dictate the composition of the bacterial community and the viability of distinct microbial niches (Rubin and Leff 2007; Mathur et al. 2007). This is particularly true of chemicals involved in alternate terminal electron accepting processes (TEAPs). In stratified lakes with high hypolimnetic oxygen demand, redox gradients spanning an array of TEAPs can develop. Different metabolic guilds can be found along these gradients in freshwater lakes (Lehours et al. 2005; Garcia et al. 2013; Salcher, Pernthaler, and Posch 2010), as is commonly observed in Winogradsky columns (Babcsányi, Meite, and Imfeld 2017; Rogan et al. 2005). Historically, our understanding of microbial distribution along spatial and temporal redox gradients has been methodologically limited. However, using recent advancements in next-generation sequencing of 16S rRNA gene amplicons, many members of highly complex microbial communities can be rapidly identified in many samples, which allows for the relative quantification of phylogenetic groups (Youngblut et al. 2013; Huse et al. 2008) Several 16S-based studies have identified compositional patterns along chemical gradients in marine (Gilbert et al. 2012; Fuhrman et al. 2006), freshwater (Jones, Newton, and McMahon 2009; Wu and Hahn 2006; Newton et al. 2007; Garcia et al. 2013), and other ecosystems. Several of these studies include investigating microbial communities along redox gradients (Garcia et al. 2013; Small et al. 2014; Rojas-Jimenez et al. 2021).

Freshwater lakes are focal points for biogeochemical cycling on the landscape and are ideal study sites for investigating the reciprocal influences between microbial communities and environmental conditions. The diversity and taxonomic composition of the freshwater microbiome has been shown to vary with environmental conditions (Newton et al. 2011; Linz et al. 2017; Paver, Newton, and Coleman 2020). Interannual variation in nutrient inputs, temperature, and a wide array of other environmental factors suggest that multi-year microbial observations are needed to adequately describe describe community dynamics(Linz et al. 2017; Shade et al. 2007). Dimitic lakes with an anoxic hypolimnion pose an especially interesting study system as the biannual mixing events cause the formation of temporary redox gradients. These gradients have been shown to facilitate the growth of a diverse microbial community (Garcia et al. 2013; Rojas-Jimenez et al. 2021). With the seasonal mixing and reforming of the redox gradient, the microbes also shape the environmental conditions as the lake stratification continues (Yu et al. 2014; Morrison et al. 2017). In addition, environmental parameters such as wind speeds, air temperature, and surface turbulence can impact a lake’s thermal stability which in turn affects the oxycline, redox gradient, and length of stratification (Kerimoglu and Rinke 2013; Foley et al. 2012). However, there is little understanding of how interannual variation in the timing of anoxia, hypolimnion temperature, and carbon loading (Ladwig et al. 2020) impact the long term changes of the microbiome in such systems (Arora-Williams et al. 2018; Garcia et al. 2013). This knowledge gap is due to the lack of multi-year studies on microbial community diversity across dynamic redox gradients, which are limited by logistical complications and long-term funding requirements.

In this study, we collected samples from a depth profile in the water column from Lake Mendota, a relatively large eutrophic dimitic lake, in 2016 and 2017. The samples were collected in the summer and fall months when the lake was stratified. During these months, the epilimnion (top layer) is oxygenated and warm whereas the hypolimnion (bottom layer) is cold, dark, and anoxic. The metalimnion, or the transition between these two zones, is characterized by a steep gradient of dissolved oxygen, temperature and sulfide. We collected samples on similar dates in 2016 and 2017 (three time points per year) to assess annual differences in the diversity and taxonomic composition of the microbial consortia across depths. While the onset of stratification and anoxia was similar, large differences in the redox status of the water column were observed between the two years. We used 16S rRNA gene amplicon sequencing to track changes in community composition both temporally and in relation to varying redox status. We found that between 2016 and 2017, many major phyla and classes were consistently detected, however, microbial groups at lower (i.e. more finely resolved) taxonomic levels showed more variance between the years. We linked the variation in several of these groups to changes in the redox gradient based on expected metabolic function of those organisms, especially near the oxic-anoxic interface. Overall, this paper shows significant chemical and microbial variation between 2016 and 2017 and highlights the importance of using multi-year, depth-discrete studies to characterize the reciprocal relationship between microbial communities and their environment along transient redox gradients.

## Results/Discussion

### Thermal stratification

While the onset of anoxia in 2016 and 2017 was comparable, there were interannual differences in the late stages of redox progression below the epilimnion between 2016 and 2017. Thermal stratification was firmly established by mid- to late May in both years (Figure S2a and b). The hypolimnetic temperature was higher in 2017 than in 2016 (11-12 °C vs 13-14 °C, respectively), likely due to warmer surface waters before stratification was established. Correspondingly, the thermocline was approximately 1-2 meters deeper in the late summer months of 2017 (~11-12 m) than 2016 (~9-10 m). The warmer hypolimnion likely also contributed to the less stable thermocline and earlier thermal destratification observed in 2017. Anoxia developed in the hypolimnion around the same time both years with dissolved oxygen (DO) reaching 0 in the deep hypolimnion by early June (Figure S2c and d). By the time we collected samples in August, the oxycline (where oxygen goes below detection) was at 11 m, near the thermocline. It then followed the downward migration of the thermocline to about 14 m in October before mixing. Thus, nearly the entire hypolimnion was at least suboxic throughout the sampling period.

### Redox status

While anoxia was first detected around the same time each year, measurements of the most prominent terminal electron acceptors (TEAs) suggest prominent differences in the redox state between years (Figure 1). We observed a large spike in nitrate/nitrite in August 2017 at 10 m, which was not seen in 2016. These peaks commonly occur in the metalimnion in July and August as ammonia from the hypolimnion is oxidized (Brock 2012) due entrainment of metalimnion or hypolimnion water. Nitrate and nitrite were then consumed through denitrification as the redox status continued to decrease. Although this nitrate/nitrite spike did not appear in our data in 2016, it is possible that it occurred during a time when we did not sample. This provides further evidence that the hypolimnion was more biogeochemically reduced in 2016 than 2017. Dissolved manganese, assumed to be in the reduced Mn(II) form, was present in similar concentrations (0 – 5 μM) throughout the water column in both years. We observed localized dissolved Mn peaks at the oxic-anoxic interface during late fall in both years which could indicate localized cycling (Oldham et al. 2017). This peak was more pronounced in 2016, suggesting elevated Mn cycling in this year, possibly due to a compressed redox gradient. The most prominent interannual difference was in the sulfide profiles. Overall, sulfide levels in 2016 were much higher than in 2017. Additionally, in September and October, 2016 had a much steeper gradient, increasing from 0 to 60 μM between 11 and 12.5 m. Conversely, in 2017, sulfide did not reach 60 μM until about 15 m and exhibited a much less steep gradient. Overall, these data again confirm that the redox status of the hypolimnetic water was substantially lower in 2016 than it was in 2017, likely indicating greater microbial activity in the hypolimnion in 2016. This is unexpected because the hypolimnion was warmer in 2017, which might be expected to drive higher microbial activity. In turn, this lower redox status generated a stronger redox gradient near the oxic-anoxic interface during late fall in 2016. This strong gradient is more likely to support redox processes that rely on such steep gradients, such as sulfide oxidation with O2 or Mn cycling (Taillefert and Gaillard 2002).

**Figure 1.**
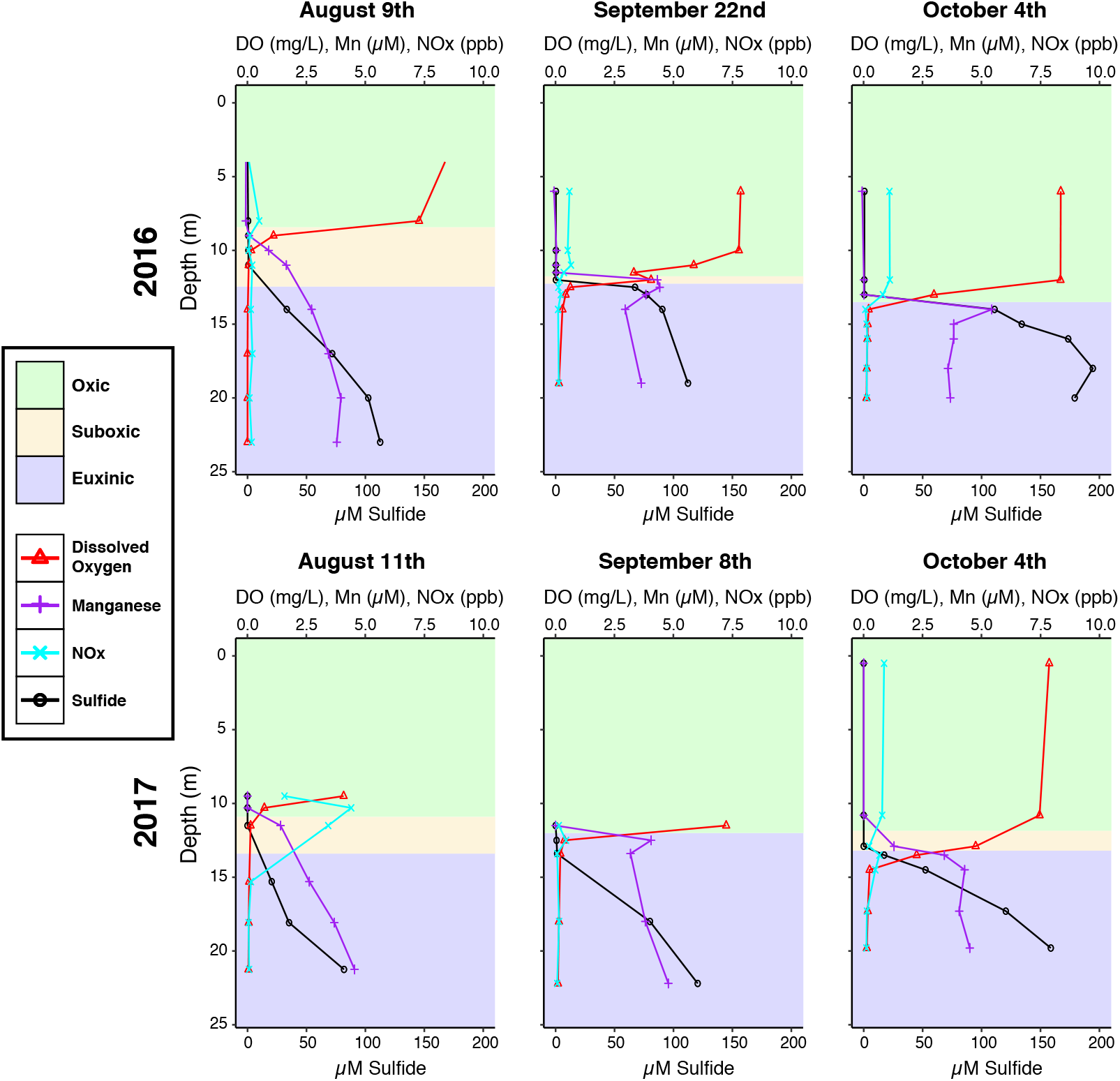
Geochemistry of Sampling Dates. Geochemical profiles of the six sampling dates. The top x axis applies to ppb Nitrate/Nitrite, μM dissolved manganese, and mg/L dissolved oxygen whereas the bottom x axis refers to μM Sulfide. Redox status is indicated by the corresponding shading color on the graph.

### Alpha diversity

We first investigated the alpha diversity of all samples included in our sequencing efforts using amplicon sequence variants (ASVs), which is the finest scale of taxonomic resolution possible using our 16S rRNA gene amplicon sequences. Previous work has shown that alpha diversity tends to be higher in the hypolimnion than in the epi- or metalimnion (Yu et al. 2014; Shade et al. 2012). We calculated the Shannon Diversity Index for each sample (Figure S1). The two-way ANOVA reveals a substantial effect of both redox state (p < 0.0001) and year (p = 0.0005) on the Shannon diversity, but no interactive effect (p = 0.56). Consistent with previous work, hypolimnetic microbial communities were more diverse than epilimnetic or metalimnetic communities. Overall, microbial communities in 2016 were more diverse. This co-occurred with the lower redox state of the hypolimnion in 2016, which led us to hypothesize that the 2016 community may contain some organisms that were reliant on sharper redox gradients in 2016.

### Beta diversity

In addition to examining diversity within each sample, we sought to identify potential drivers of community composition change by observing the relationship between changing redox conditions and microbial community composition shifts. An NMDS ordination based on Bray-Curtis dissimilarity revealed separation of the samples based on both redox status and year (figure 2). To test these observations statistically, we ran a UniFrac distance-based PERMANOVA on our samples, which showed that clustering by year was significantly different (R^2^ = 0.036). On the other hand, clustering by redox status was not significantly different (R^2^ = 0.140), likely due to the overlap between the suboxic and euxinic samples, especially in 2016. These results are consistent with our NMDS ordination where there is greater separation by year than by redox state. Because the redox status of the water plays a major role in microbial metabolism by controlling TEA availability, we would expect major differences in the community composition under these different conditions. In 2016, we observed a prominent separation between the oxic samples and the other two classifications, but substantial overlap between the suboxic group and the euxinic group. In 2017, the oxic samples were still distinct, but there was also little overlap between the suboxic and euxinic groups. This suggests that suboxic and euxinic microbial communities were less similar to each other in 2017 than 2016. Considering that our alpha diversity analysis shows that suboxic waters of 2017 were less diverse, it is possible that some organisms that are viable in both euxinic and suboxic waters were not viable in the suboxic waters of 2017, possibly due to differences in redox status. While these differences in redox status were noticeable, overall the redox progression followed a similar pattern. However, this is not observed in the ordinations, as there is little overlap between the two years, suggesting significant differences between the microbial populations in each year (Figure 2). This is true for samples within all redox classifications.

**Figure 2.**
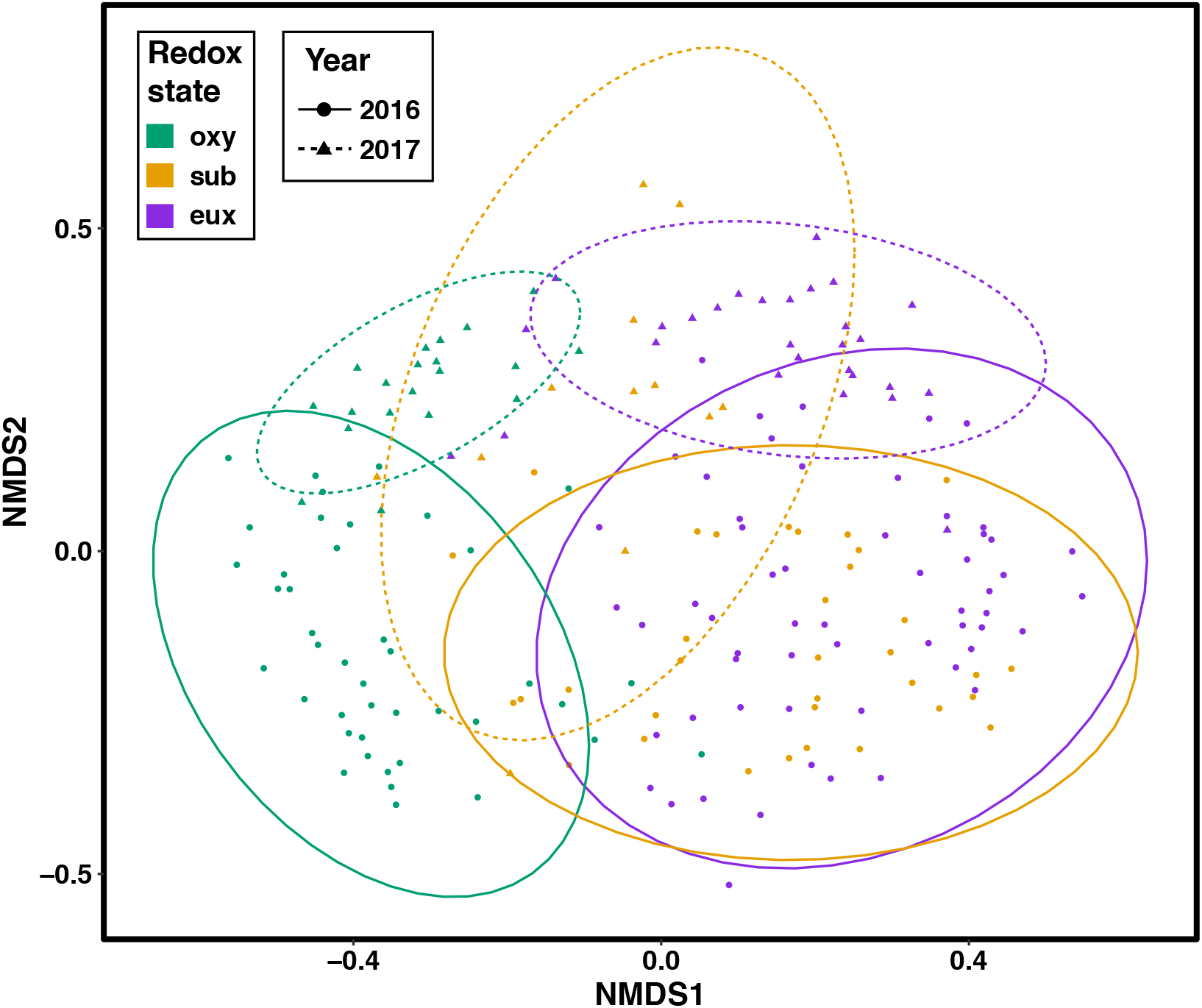
ASV variability by year and redox state implies temporal distance explains more variability than spatial distance. Nonmetric Multidimensional Scaling (NMDS) using Bray-Curtis distance to display dissimilarity between each sample. Dissimilarity was calculated using all ASVs as a distance matrix. Ellipses represent the clustering of samples by redox state (color) and year (line type) using a 95% confidence level. ASVs between oxy and eux have little to no overlap both years, whereas both eux and oxy have substantial overlap with sub samples. Within each redox state, 2016 and 2017 do not have much overlap.

We next investigated beta diversity patterns within subsets of the dataset to further probe the extent of interannual variation. Specifically, we removed the oxic samples (DO > 1 mg/L) from the analysis to focus on the relationship between community composition and alternative TEAs. First, we specifically wanted to investigate the relative impact of the hypolimnetic redox status on community composition and how this related to interannual variation. A canonical correspondence analysis (CCA) ordination is useful for incorporating environmental parameters into an analysis of beta diversity. In our CCA ordination of suboxic and euxinic samples, samples were strongly separated by year (Figure 3A). There was also strong separation of the samples by redox status. Samples separated more strongly by year than by redox gradient, but the redox status is more of a continuous variable, whereas year is discrete, likely contributing to this difference. Although at first this may seem contradictory to our PERMANOVA and NMDS ordination where there was no significant separation by redox status, dissolved Mn and sulfide are used as constraining variables in this CCA. These two variables varied by year, and especially so in suboxic and euxinic depths. Having separation of suboxic and euxinic samples along Mn and sulfide emphasizes temporal distance as a driver for community diversity because of the volatile nature of Mn and sulfide with respect to time.

**Figure 3.**
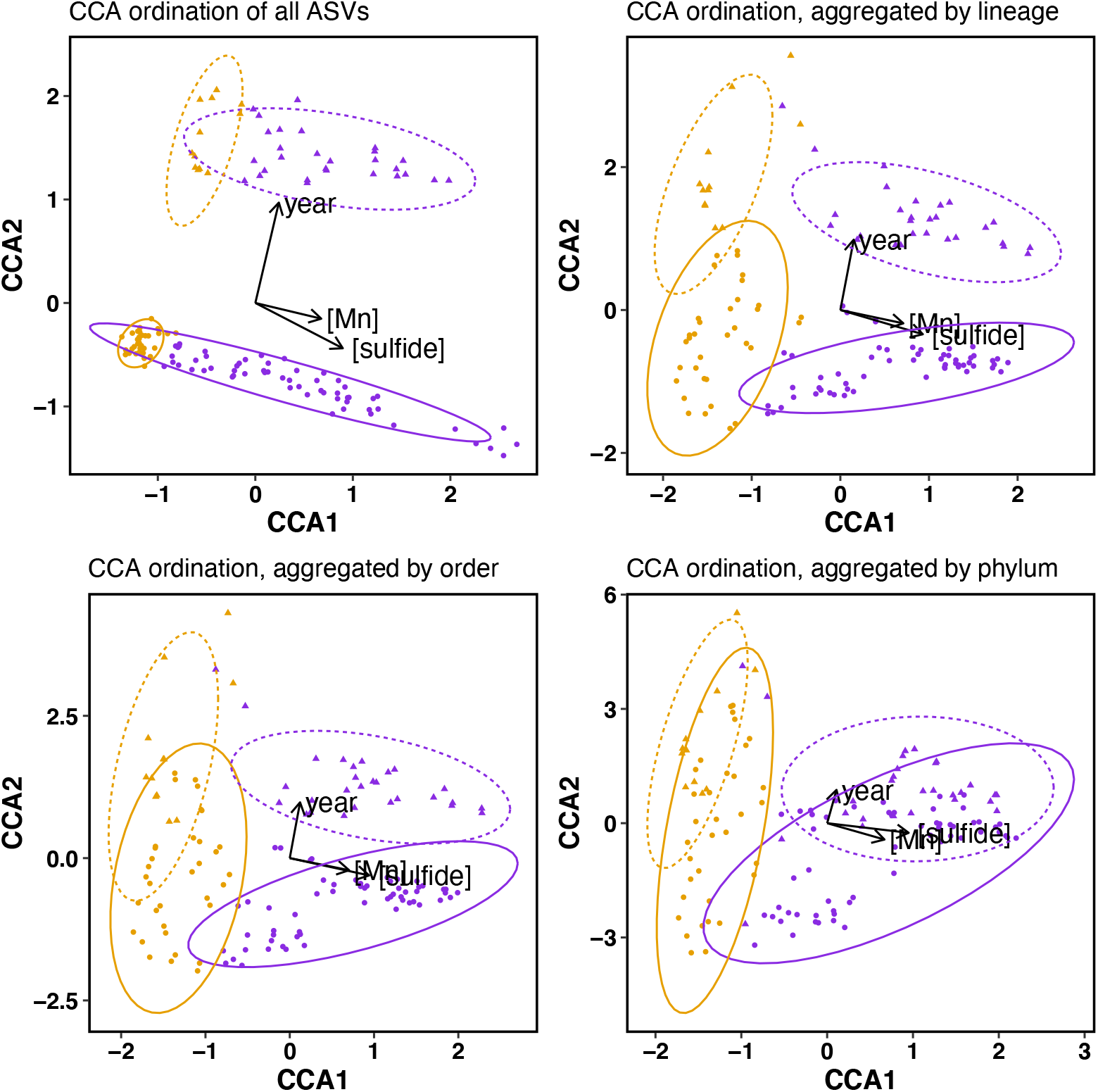
Constrained ASV Variability of Anoxic Samples of Different Taxonomic Levels. Canonical correspondence analysis (CCA) on anoxic samples using Bray-Curtis distance to display dissimilarity between the normalized ASVs of each sample. Sulfide and manganese levels and year are used as constraining variables. Ellipses represent the clustering of samples by redox state (color) and year (line type) using a 95% confidence level. Sub samples are yellow, Eux samples are purple, 2017 has triangle points with a dashed line ellipse and 2016 has circle points with a solid line ellipse. Unclassified phylum, orders and lineages are not included in this analysis. There is strong separation of the samples by year when all ASVs are considered. Suboxic and euxinix samples have small overlap with the water chemistry vectors point towards the euxinic samples. Within each redox state, 2016 and 2017 have substantial overlap. In the phyla ordination, each redox state has overlap between the two years, however this overlap diverges in the order and lineage ordinations. In the lineage ordination, euxinic between the two years has no overlap.

The extent of the variation in community composition attributed to interannual differences as opposed to the redox gradient is surprising but does not necessarily indicate strong functional differences. While 16S rRNA gene based identification cannot consistently and reliably predict most metabolic functions, it is likely that, on average, functional guilds are more closely related. We thus hypothesized that interannual differences in the microbial community would be less pronounced at coarser taxonomic levels. To visualize this, we performed the CCA ordination on data that had been aggregated by family, order, or phylum. We observed decreasing interannual variation as the data was aggregated by higher taxonomic levels (Figure 3B-C). There was no difference in the impact of the redox parameters on sample separation at any of the taxonomic levels. This suggests that community composition differences across years are more pronounced when taxa are split into finer and finer groups due to stochastic processes that determine which strains or closely related species within a functional guild assemble, rather than substitution of whole functional guilds across years. More simply, the same functional guilds are present each year but different species or strains comprise those guilds. One important note is that CCA analyses are sensitive to rare species. This bias means that if a sample has an organism that is not present in other samples, it will be weighed more in its overall difference from other samples (Ramette 2007). The fact that the groups converge but stay cohesive within themselves (ellipses do not change shape significantly at finer taxonomic levels) despite this bias emphasizes that the contrast is not excessively driven by aggregation of rare taxa.

Overall, our ordinations and PERMANOVA suggest that there is substantial year-to-year variation in the community composition, but that this variation emerges mainly at finer taxonomic scales. At the phylum level, there is little variation in the microbial community year to year within the different redox layers. On the other hand, the variation related to redox gradients is robust over all taxonomic levels we investigated, suggesting a more consistent functional response of the microbial community to redox gradients, as might be expected based on fundamental principles of environmental microbiology (cite Brock textbook!). These results highlight the need to study freshwater systems over multiple years to fully comprehend microbial community patterns in spatially-resolved datasets. Future work on intra- and inter-annual differences in microbial community function would help resolve some of these questions, such as through functional assays or shotgun metagenomic sequencing to examine differences in functional gene abundance.

### Taxonomic analysis

To further investigate these taxonomic shifts in the microbial community underlying the above trends in beta diversity, we examined the taxonomic composition of the microbial community from a subset of samples. We selected three profiles from each of the two years to investigate in detail, chosen to match up by date. We assigned taxonomic classifications to each ASV using a custom freshwater 16S rRNA gene database and investigated the spatial and temporal variation in the microbial community composition at different taxonomic levels.

### Taxonomy - phyla

The most coarse taxonomic level, phylum, was a natural starting point to visualize the most general trends in the microbiome. The relative abundance of phyla has been shown to change dramatically with depth in several other depth-discrete sampling studies (Tran et al. 2021; Rojas-Jimenez et al. 2021; Baatar et al. 2016; Garcia et al. 2013). These changes in phyla are often attributed to the redox gradients throughout the water column (Jones, Newton, and McMahon 2009; Wu and Hahn 2006; Newton et al. 2007; Garcia et al. 2013). However, less is known about interannual differences in taxonomic groups across these re-occurring redox gradients. Based on our ordinations (Figure 4), we expected to see greater differences in phylum-level community composition along redox gradient than between the two years. We observed substantial variation in the phyla across the redox gradients, but very little interannual variation, as we would expect from the ordinations. In the epilimnion, Actinobacteria and Cyanobacteria are the most prevalent phyla, usually consisting of about 30% relative read abundance each in the epilimnion, but dropping to about 10% and 5% in the hypolimnion, respectively (Figure 4). The presence of even a small number of Cyanobacteria at the anoxic bottom of the lake is surprising considering their association with aerobic metabolism. However, this has been observed elsewhere (Tran et al. 2021). Actinobacteria are known to include the cosmopolitan epilimnetic lineage acI (order Nanopelagicales), and account for much of the abundance in the epilimnion. Kiritimatiellaeota, on the other hand, are absent in the epilimnion but become one of the more common phyla in the hypolimnion at around 15% relative abundance. Kiritimatiellaeota have been previously shown to be highly abundant in the hypolimnion of Lake Mendota, are known to thrive in anoxic conditions, and degrade polysaccharides (Spring et al. 2016; Peterson, McDaniel, Schmidt, Lepak, Tran, et al. 2020). Firmicutes were also primarily abundant in the hypolimnion (10-15%), which is consistent with their documented role as fermentative organisms (Weber, Achenbach, and Coates 2006). Other phyla are consistently abundant throughout the water column. For example, Proteobacteria are the most prevalent phyla in the hypolimnion usually consisting of about ~40% relative read abundance, but also are present at 20% in the epilimnion. This is likely due to the high functional diversity of this phylum, which is confirmed by the variation at finer taxonomic levels across the redox gradients (see below). The abundances of each of these phyla show very little interannual variation. However, there are some lower-abundance phyla that do show interannual variation. For example, the candidate WS4 phylum is only present in the hypolimnion in early August of 2016. Armatimonadetes was also only present in the hypolimnion of 2016, but was there in August, September and October. Organisms such as these might be reliant on the lower redox status that was present in 2016. Alternatively, their presence in 2016 but not 2017 could be strictly due to stochastic processes. Regardless, their appearance likely contributes to the greater richness observed in 2016 relative to 2017 (see alpha diversity, Figure S1).

**Figure 4.**
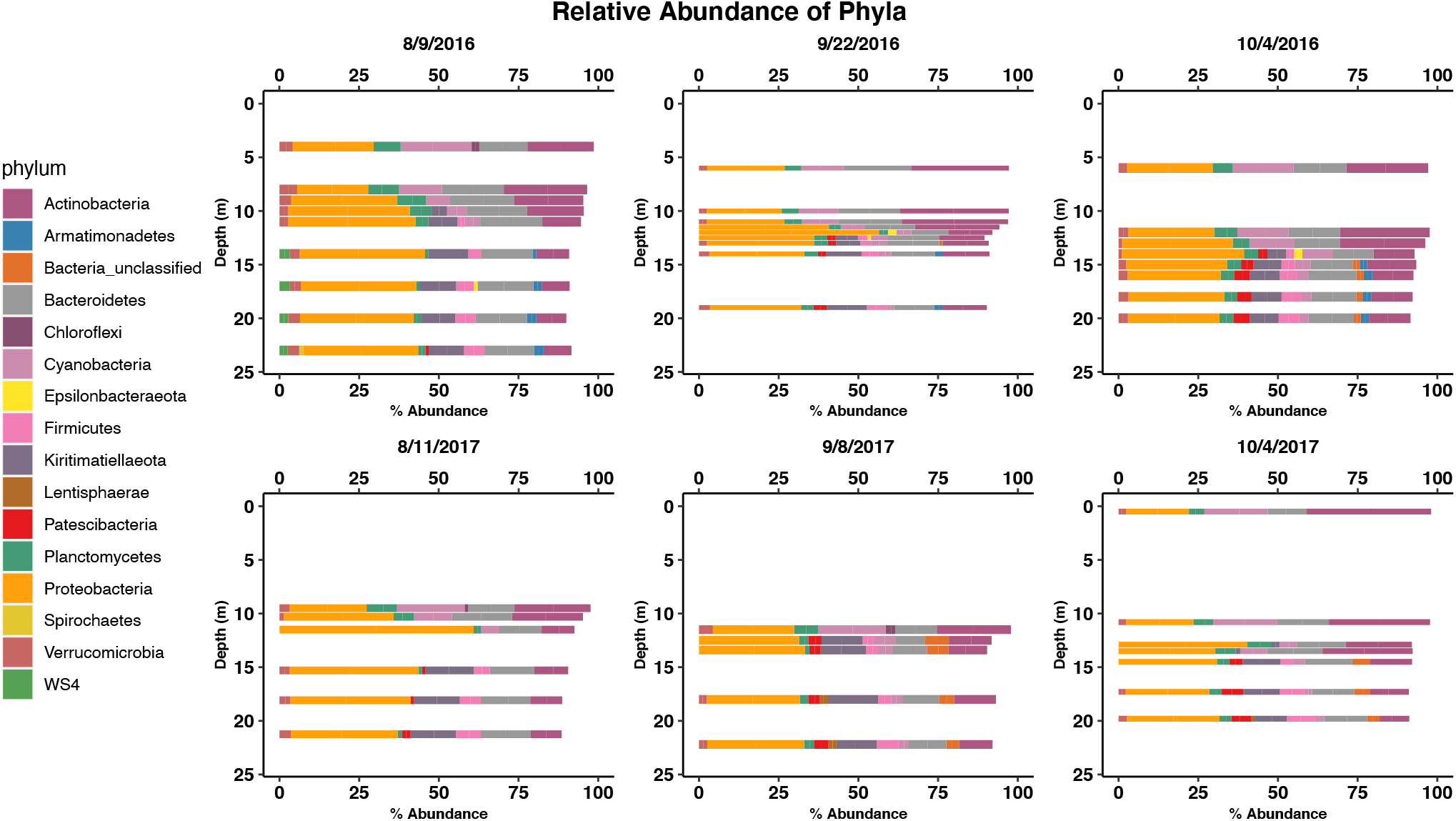
Vertical Distribution of Phyla. Bar charts displaying the relative abundance profiles of bacterial phyla from three sampling dates in 2016 and 2017. Samples for each date were taken at several discrete depths above, at, and below the oxycline. Bacterial communities from the samples are classified at the phylum level and phyla accounting for less than 1% of the overall read abundance are not shown.

### Taxonomy – class, order, family

We next investigated the abundances of several finer-scale taxonomic groups. We selected a subset of taxonomic groups to highlight the observed trends and show how the changing redox status can affect the microbial community.

We first investigated Proteobacteria composition at lower taxonomic levels since they were one of the most prevalent phyla in both oxic and anoxic waters. At the class level, Proteobacteria in our samples exclusively consisted of Gammaproteobacteria, Alphaproteobacteria and Deltaproteobacteria (Figure 5). Within Proteobacteria, Gammaproteobacteria are the most abundant in the epilimnion, however their abundance sharply declines in the hypolimnion where Deltaproteobacteria become more prevalent. Here we note that our general reference database for classification was SILVA XXX, which incorporates a re-organized Bacterial phylogeny that most significantly moved the old class Betaproteobacteria as an order into the Gammaproteobacteria class. Consistent with literature showing Betaproteobacteria to be some of the most common bacteria in upper waters of lakes (Newton et al. 2011), Betaproteobacteria accounted for a majority of Gammaproteobacteria in the epilimnion (Figure S3). The markedly higher relative read abundance of Deltaproteobacteria in the hypolimnion is parallel with a general increase of sulfide in the hypolimnion and a shift towards Desulfobacterales, which make up 40% to 70% of the Deltaproteobacteria reads (Figure 5). Considering that Desulfobacterales are usually associated with sulfate-reduction it is not surprising they are most abundant in euxinic waters (Figure S4). Another order of Deltaproteobacteria, Desulfuromonadales, are dominated by Geobacteraceae, specifically *Geobacter* at the family and genus level. These microbes were most abundant at the oxic/anoxic interface, where they increased in read abundance with depth almost ten-fold before receding in the hypolimnion to about 0.1% relative abundance (Figure 6). This corresponds with a striking increase in dissolved Mn, at the depth where we suspect elevated Mn cycling occurs, suggesting these organisms could be respiring Mn. *Geobacter* are well-known Mn reducers (Lovley and Phillips 1988), and we recently identified several Geobacterales genomes in Lake Mendota, all of which contained a porin-cytochrome c complex (PCC) operon, further supporting this hypothesis (Peterson, McDaniel, Schmidt, Lepak, Janssen, et al. 2020). In our data, Desulfuromondales are about 3-4 times more abundant in September and October 2016 than they were one year later, which coincides with a more prominent metalimnetic Mn maximum in 2016 than in 2017. A previous study also demonstrated *Geobacter*-associated metal reduction genes spiking at similar depths in a lake that is of nearly identical depth in 2013 (Arora-Williams et al. 2018). This difference in *Geobacter* abundance highlights the importance of interannual variability in redox status in determining the abundance of select metabolic guilds.

**Figure 5.**
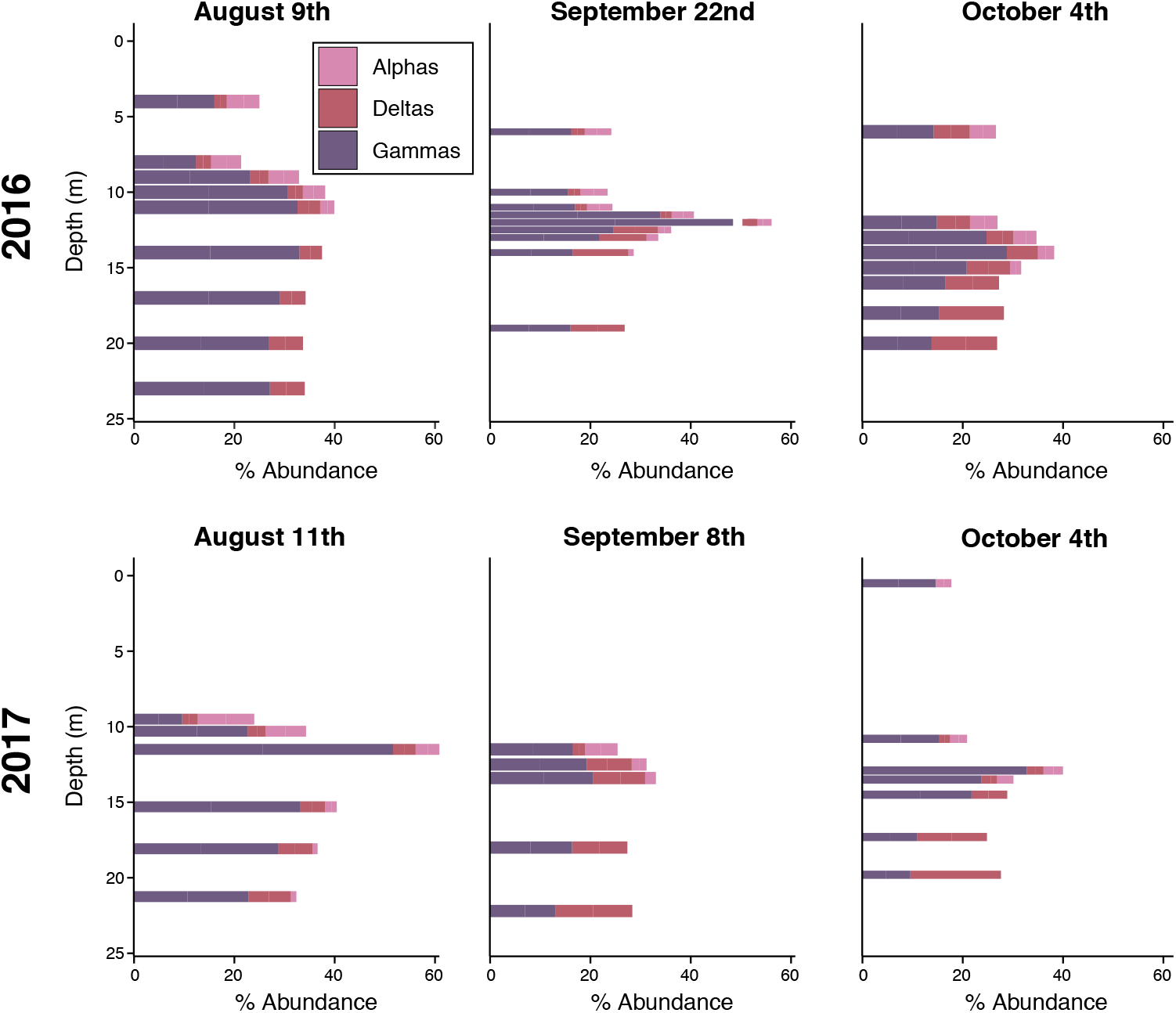
Composition of phylum Proteobacteria. Bar charts displaying the distribution and the composition of classes in phylum Proteobacteria for each sample. Samples for each date were taking at several discreet depths above, at and below the oxycline. Proteobacteria composition changes throughout the water column with respect to the chemistry of the water and the classes’ metabolic tendencies. Gammaproteobacteria tend to dominate in the epilimnion while deltaproteobacteria greatly increase in relative abundance in the hypoliminion.

**Figure 6.**
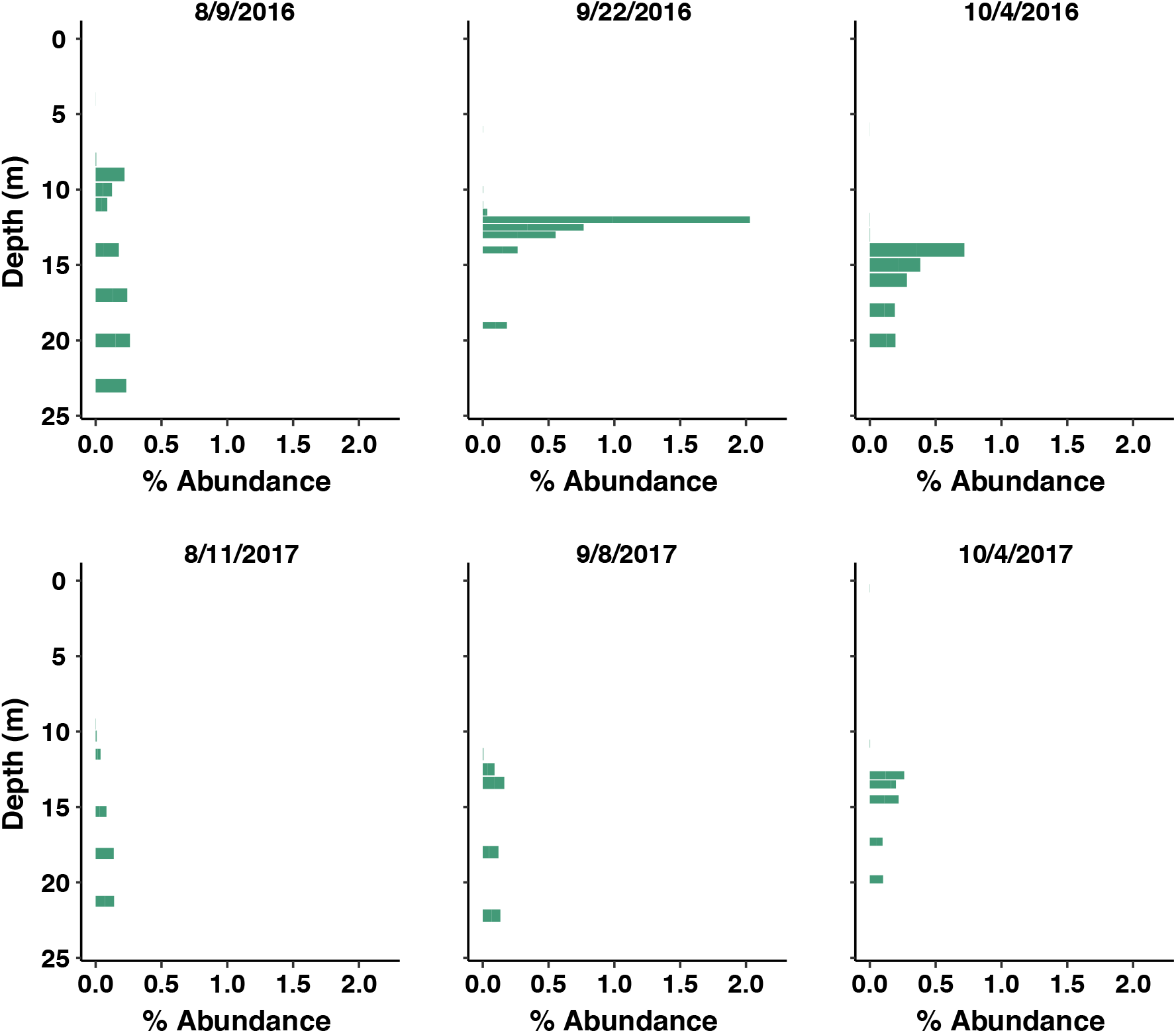
Vertical Distribution of Desulfuromonadales. Bar charts displaying the distribution and relative abundance of order Desulfuromonadales. Samples for each date were taking at several discrete depths above, at and below the oxycline. This graph shows the relative abundance of order Desulfuromonadales with respect to all ASVs. Desulfuromonadales spike at the oxic-anoxic interface. In our dataset, Desulfuromonadales consist entirely of Geobacteraceae and *Geobacter* at the family and genus level respectively.

We also further investigated the taxonomic composition of the Cyanobacteria, particularly to identify the lineages living in the hypolimnion. Despite their reputation as being photoautotrophic, Cyanobacteria have been known to grow in dark and anoxic waters (Callieri et al. 2019; Tran et al. 2021). In our samples, at the class level, Cyanobacteria were almost exclusively Oxyphotobacteria (~97%). The main orders of Oxyphotobacteria include Synechococcales (~15%-80%), Phormidesmiales (~5%-25%) and Nostocales (~5%-50%) (Figure S5). Despite the large range in relative read abundance of some of these orders, for the most part Synechococcales are the dominant order, especially in 2016. Synechococcales become a larger portion of cyanobacteria in the hypolimnion in 2016, however Synechococcales are evenly distributed in 2017. Nostocales and Phormidesmiales are evenly distributed in 2017, but wane in abundance in the hypolimnion in 2016. Nostocales are more abundant throughout the entire water column in 2017 than 2016. There have been some studies that have reported that hypolimnetic photosynthetic picoplankton communities become dominated by phycoerythrin-containing Synechococcus during seasonal thermal stratification (Becker, Richl, and Ernst 2007; Fahnenstiel and Carrick 1992). This could explain why in August and September 2016, Synechococcales became so dominant, however the same level of abundance was not seen one year later. Although the exact mechanism by which the cyanobacteria population varies in 2016 versus 2017 cannot exactly be elucidated by our study, Cyanobacteria variation is nevertheless intriguing due to the phylum’s abundance and role as a primary producer in the ecosystem. Future work should also focus on whether cyanobacterial cells in the hypolimnion are active or simply sinking while senescing.

Members of other phyla, such as the Kiritimatiellaeota phylum, have limited fine-scale taxonomic resolution, likely due to poor representation of this phylum in databases. This complicates comparisons of closely related lineages between years. All ASVs classified as Kiritimatiellaeota are Kiritimatiellae at the class level and mostly (95%+) WCHB1-41 at the order level. However, they are mostly unclassified (95%+) at the family, genus, and species levels. This limited classification is likely due to the novelty of the phylum, which was only discovered recently (Spring et al. 2016). Kiritimatiellaeota increase in abundance with time in the hypolimnion in Mendota over both years of study, however they are present in some epilimnion samples. This may suggest that there are aerobic lineages of WCHB1-41 that have yet to be described. Although not much are known about this lineage, Kirimatiellaeota are mostly found in anoxic waters, and some are known to degrade polysaccharides, so these two aspects may very well be true of WCHB1-41 (Van Vliet et al. 2019).

A significant difference in the microbial community composition of the water column between 2016 and 2017 is in the key anaerobic photosynthetic groups class Chlorobia and family Chromatiaceae. Chlorobia are one of the two major groups of green sulfur bacteria (GSB) and are photolithotrophs and obligate anaerobes. Chromatiaceae are a subset of Gammaproteobacteria commonly known as purple sulfur bacteria. Both groups were observed in Lake Mendota only during September and October 2016 at the oxic/anoxic interface, where sulfide drastically increased with depth. Specifically, Chlorobia increased to around 0.1% of all ASVs and Chromatiaceae increased to about 0.25% of all ASVs (Figure 7). Anaerobic photosynthesis often uses sulfide as a terminal electron acceptor to fix carbon, so this process is relevant in the sulfide and carbon cycles. Due to anaerobic photosynthesis requiring light, high sulfide, and anoxia, these types of organisms are often found in stagnant water bodies and microbial mats (Van Gemerden and Mas 1995). The required conditions for anaerobic photosynthesis are not easily met in a deep dimictic and highly eutrophic lake like Mendota. Intense cyanobacterial blooms in the upper water column are thought to shade the depths below, excluding other phototrophs. However, anoxygenic phototrophs can be active under extremely low-light conditions (cite Brock textbook?). Furthermore, Chromatiaceae also include chemolithotrophs that can oxidize sulfide using oxygen (though Chlorobia are usually thought to be strictly photolithotrophic) (cite Brock textbook). The redox conditions present at the oxic-anoxic interface should be able to support such chemolithotrophic growth. On September 9^th^, 2016 at 12.5 m (the first euxinic depth), D.O. drops to 0.625 mg/L and sulfide jumps to 67 μM and this produces the largest spike in abundance of the groups discussed. These conditions continue through to October 4^th^, 2016, where at 15 m we observed a similar low oxygen/high sulfide environment. September and October 2017 do not have any shallow euxinic depths with overlap in oxygen and sulfide due to the redox gradient not being as strong that year. Thus, it makes sense that these types of microbes were not able to make a living under the conditions observed in 2017. Chlorobia in previous studies have shown spatial abundance variability positively correlated with NH_4_-N so the differences in the redox gradient between 2016 and 2017 may very well be explanatory for our temporal variability (Edberg, Andersson, and Holmström 2012; Eraqi et al. 2021). The temporal disparity of microbes capable of anaerobic photosynthesis suggests that certain functional guilds involved in important environmental cycles will not appear consistently every year.

**Figure 7.**
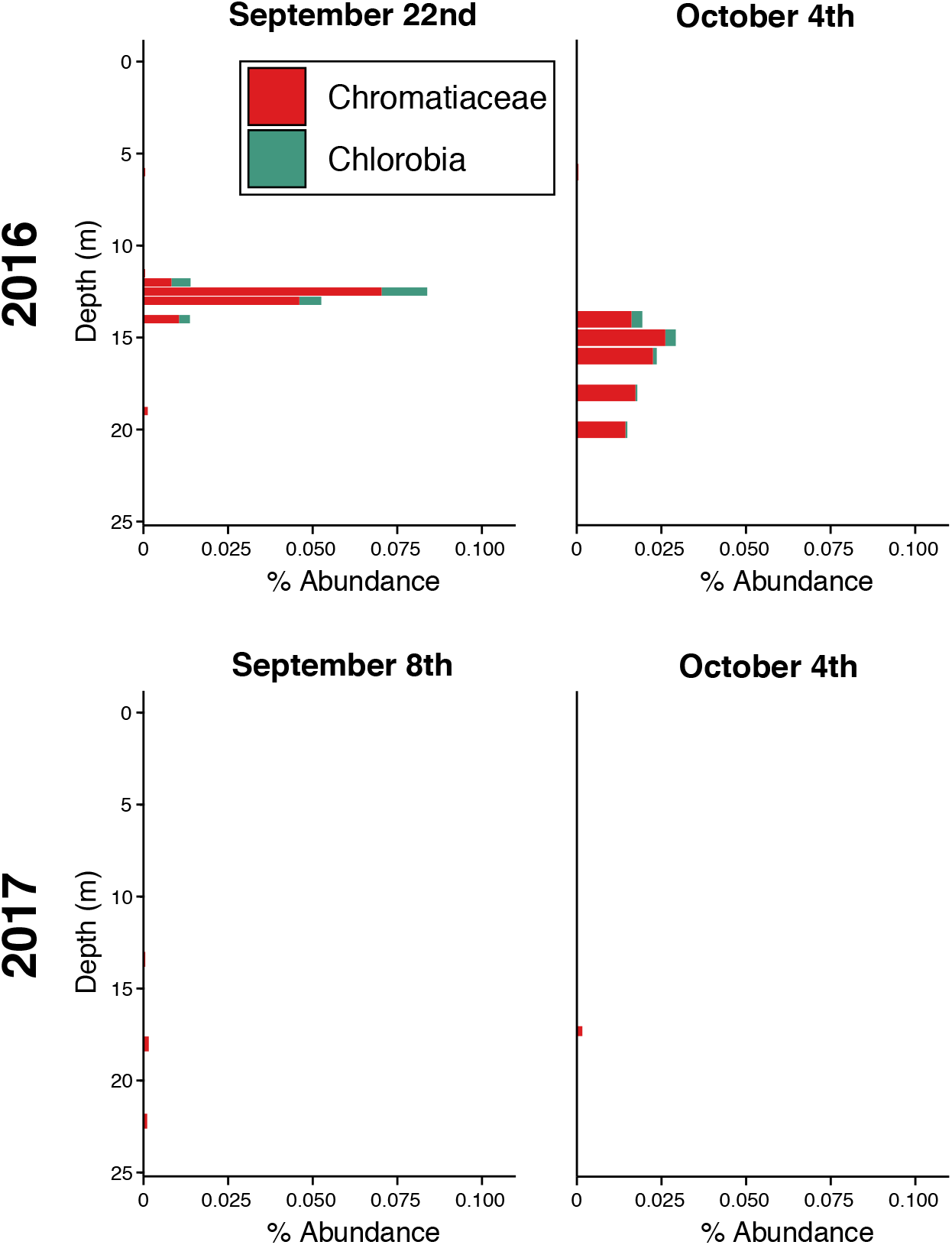
Vertical Distribution of lineage Chromatiaceae and class Chlorobia. Bar charts displaying the distribution and relative abundance of lineage Chromatiaceae and class Chlorobia. Red is Chromatiaceae and Brown is Chlorobia. Samples for each date were taking at several discrete depths above, at and below the oxycline. This graph shows the relative abundance of lineage Chromatiaceae with respect to all ASVs. These two lineage and class are prevalent in anoxic photosynthetic and are mostly only seen in 2016.

Ignavibacteria, previously considered to represent a class within the phylum Chlorobi (now renamed to be Bacteroidota) were also much more prevalent in 2016. These organisms are thought to be metabolically diverse heterotrophs incapable of photosynthesis (Liu et al. 2012). Ignavibacteria usually spiked (2.5-5% of all ASVs) when D.O. dropped below 3 mg/L and just before dissolved Mn started to sharply increase (Figure S5).

All in all, the main differences at finer taxonomic levels were found in more specialized lifestyles corresponding to a slightly different redox status, usually at the oxic-anoxic interface, between the years. For example, the anoxic photoautotrophs were seen only in 2016 and Desulfuromondales only peaked at the oxic-anoxic interface in 2016. These examples demonstrate that the presence of certain groups of microbes, some of them corresponding to unique functional guilds, can be dependent on environmental conditions that change annually. For example, it is likely that Mn reduction and lithotrophic sulfide oxidation were important biogeochemical processes in the metalimnion in 2016, but may have played a smaller role in 2017. These groups are in the minority, however, as most taxonomic groups that we investigated above the family level exhibited similar patterns from year to year.

As lakes are hot spots for geochemical cycling in the landscape, understanding their dynamics is critical to environmental research at large. As climate change, land-use change, and other outside events change lake chemistry all over the world, the need to understand how the microbiome could respond to these events is critical now more than ever(Adrian et al. 2009; Fournier, Lovejoy, and Vincent 2021). The data we have collected adds to a growing body of evidence that extrapolation from a limited number of observations in space or time provides an incomplete picture of microbial processes at larger/longer scales. How often, and for how long, do we need to sample a lake in order to understand its microbiome? Dynamic geochemical conditions in spatially structured systems such as stratified lakes make this question even more pressing. More multi-year, depth-discrete studies including detailed geochemical characterization are needed to comprehend long-term trends in lake microbial community dynamics. These types of studies will ultimately help elucidate the reciprocal relationships between microbial taxa and their environment, with implications for ecosystem-scale prediction.

## Materials and Methods

### Sample Collection

We sampled Lake Mendota, a eutrophic dimictic lake located in Madison, WI, USA. Samples were taken near the deepest part of Lake Mendota (GPS coordinates: 43.0989, −89.405). Sampling events occurred approximately bimonthly throughout the summer and fall (June - November) of 2016 and 2017. This temporal range covers early stratification and turnover of the water column. Profiles of temperature and dissolved oxygen were collected continuously with a YSI Exo2 multiparameter sonde (YSI Incorporated, Yellow Springs, OH). Temperature profiles were supplemented with temperature readings from the North Temperate Lakes Long-Term Ecological Research (NTL-LTER) station (Magnuson, Carpenter, and Stanley 2010). Dissolved oxygen profiles were supplemented with profiles collected by the Microbial Observatory NTL-LTER. All samples were collected using a peristaltic pump with an acid-washed Teflon sampling tube. Water samples for sulfide analysis were preserved in an acid-washed Nalgene (Nalgene Nunc International Corporation, Rochester, NY) HDPE plastic bottle with 1% zinc acetate. Water samples for dissolved manganese analysis were filtered with a 0.45 μm PES Acrodisc filter and immediately preserved to 1% HCl. Microbial samples for DNA extraction were collected on 0.22μm pore-size PES filters, then immediately flash-frozen on liquid nitrogen and stored at −80°C.

### Geochemical Analysis

Sulfide quantification was done using the Cline method with spectrophotometry (Cline 1969). Manganese was quantified using inductively coupled plasma optical emission spectroscopy (ICP-OES) on a Varian Vista-MPX CCD ICP-OES. Nitrate/nitrite was quantified using colorimetric spectrophotometry on an Astoria II segmented flow autoanalyzer (Astoria-Pacific). DNA was extracted from the filters via enzymatic and physical lysis preceding phenol-chloroform extraction and purification by isopropanol precipitation. Details of the DNA extraction protocol and other methods can be found in Peterson et. al. 2020. The geochemical data was collectively used to assign a redox status to each sample we collected. Samples with over 1 mg/L DO were marked as oxic samples (oxy). Samples with less than 1 mg/L DO and less than 2 μM of sulfide were classified as suboxic (sub), while those with over 2 μM of sulfide were euxinic (eux). No samples were observed with both > 1 mg/L DO and > 2 μM sulfide.

### 16S amplicon sequencing and bioinformatics

A total of 194 samples (97 locations with biological duplicates) were sequenced at the Biotech Center at the University of Wisconsin - Madison. The v3v4 region was amplified and sequenced on an Illuminia MiSeq to generate 250bp paired-end reads. Read counts for each sample ranged from 36,112 to 87,069 with a mean read count of 63,343 (Table S1/Github). The mothur SOP was followed for sequence processing, trimming, and quality control (Kozich et al. 2013). Briefly, paired end reads were trimmed and merged, then pre-clustered using a 4-nucleotide difference cut-off. Then, chimeras were removed using UCHIME (Edgar et al. 2011), resulting in 269,921 sequences with an average length of 459. These sequences are described here as Amplicon Sequence Variants (ASVs) (Callahan, McMurdie, and Holmes 2017). ASVs matching a sequences in the FreshTrain database with a percent identity greater than 98% based on blastn were classified in that database, and the remaining sequences were classified in the Greengenes database (Camacho et al. 2009; Newton et al. 2011; Rohwer et al. 2018; McDonald et al. 2012).

### Statistical methods

All statistical analysis, ordinations and taxonomic analysis were done in R version 3.6.2 (R Core Team 2019). ASVs only recorded once across all samples were removed. The ASV counts were then normalized to the total counts within every sample. Shannon Diversity Index was calculated for each sample using the diversity() function on the normalized ASVs in “vegan” (Oksanen et al. 2019). For the Nonmetric Multidimensional Scaling (NMDS) ordination a dissimilarity matrix was generated based on Bray-Curtis dissimilarity of the samples using the normalized ASVs. NMDS ordinations were performed with the metaMDS() function in “vegan”. A canonical correspondence analysis (CCA) ordination of the anoxic samples using Bray-Curtis dissimilarity was created using sulfide, manganese, and year of the sample as constraining variables. This was also done after grouping ASV counts at the phylum, class, and lineage level in each sample. Taxa that were unclassified at any taxonomic level down to the level of clustering were removed for this second analysis. The fraction of reads removed at each taxonomic level are shown in Figure S7. The CCA ordinations were implemented using the cca() function in “vegan”. Packages “dplyr” (Wickham et al. 2019) and “tidyr” (Wickham and Henry 2019) were used for data formatting and all plots were generated using “ggplot2” (Wickham 2016). Weighted UniFrac distance was used to determine similarity between samples for permutational multivariate analysis of variance (PERMANOVA). Weighted UniFrac distance was implemented using the phyloseq() function in the “phyloseq” package (McMurdie and Holmes 2013). A phylogenetic tree for the UniFrac was generated using neighbor-joining using the DistanceTreeConstructor() function in Biopython library (Cock et al. 2009) in Python 3.8.3 (Python Software Foundation, https://www.python.org/). All analyses using the weighted UniFrac distance matrix were done using only ASVs that appear more than 500 times across all 194 samples. Even with this cutoff, NMDS ordinations are impacted little (Figure S8), so this subset should still be a good representation of the samples. All scripts used for analysis are available on GitHub: (https://github.com/robertmarick/Mendota20162017). Processed 16S data, geochemical measurements, and dissolved oxygen profiles from the NTL-LTER Microbial Observatory are available on OSF: https://osf.io/p92gn/

## Acknowledgments

We acknowledge the North Temperate Lakes Long Term Ecological Research (NTL-LTER) site, Lake Mendota Microbial Observatory field crews, and University of Wisconsin - Madison Center for Limnology for the field and logistical support. Funding for geochemical analysis and 16S sequencing was provided by the Wisconsin Sea Grant College Program Project #HCE-22. We thank graduate students Tylor Rosera, Stephanie Berg, and Marissa Kneer for sampling assistance. We also thank undergraduate researchers Mykala Sobieck for sampling design and sampling assistance and Anna Schmidt, Diana Mendez, and Ariel Sorg for sampling assistance. Geochemical analyses were performed at the Water Science and Engineering Laboratory at the University of Wisconsin-Madison. Computational work was performed in part using the Wisconsin Energy Institute computing cluster, which is supported by the Great Lakes Bioenergy Research Center as part of the U.S. Department of Energy Office of Science.

## Supplementary Material

**Figure S1.**
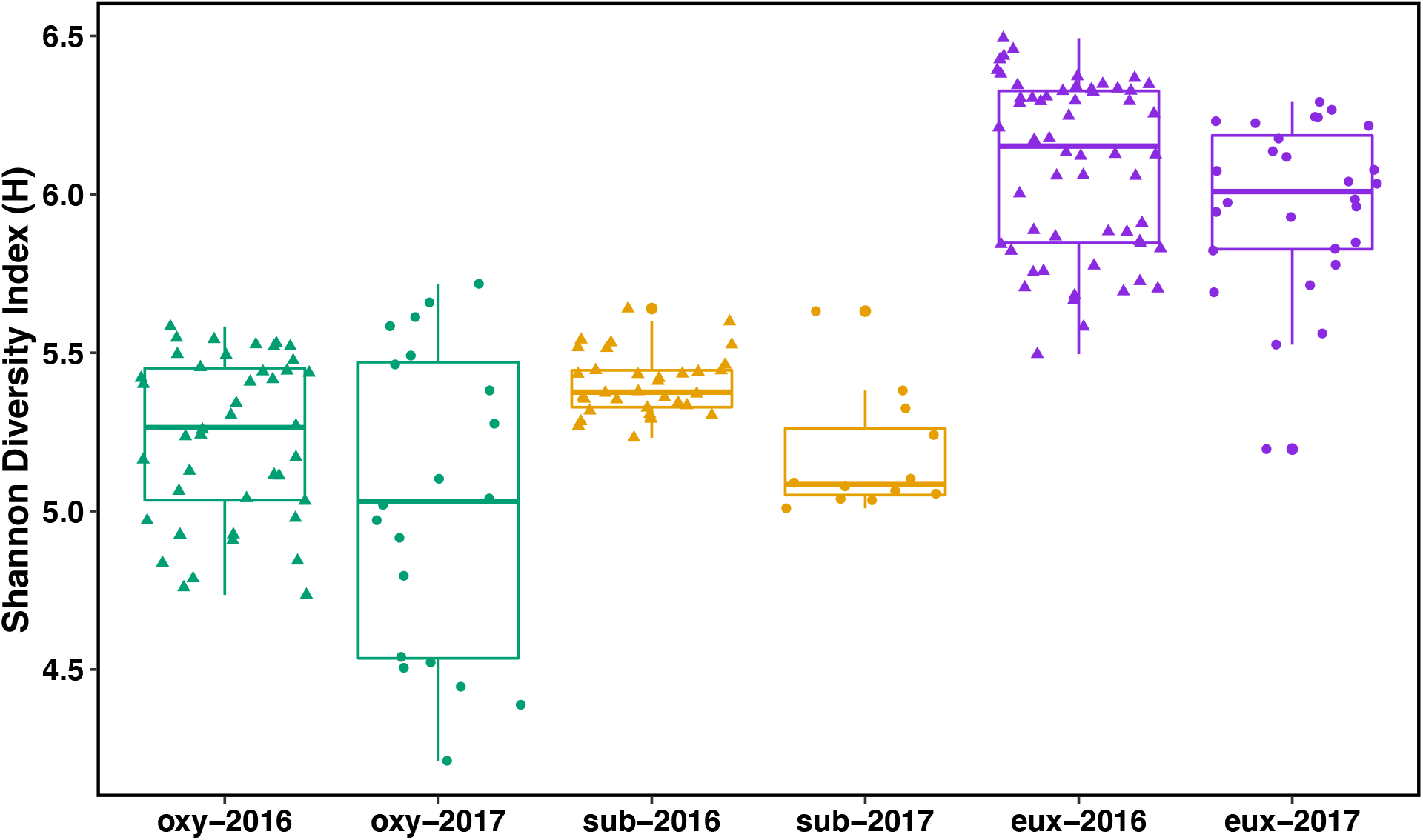
Shannon Diversity Index of All Samples. The Shannon diversity index of all samples was calculated based on a sample-by-ASV matrix and plotted according to the sample’s group. A two-way ANOVA showed significant difference between year (p = 4.61e-4) and redox status (p < 2e-16) but no significant interactive effect.

**Figure S2.**
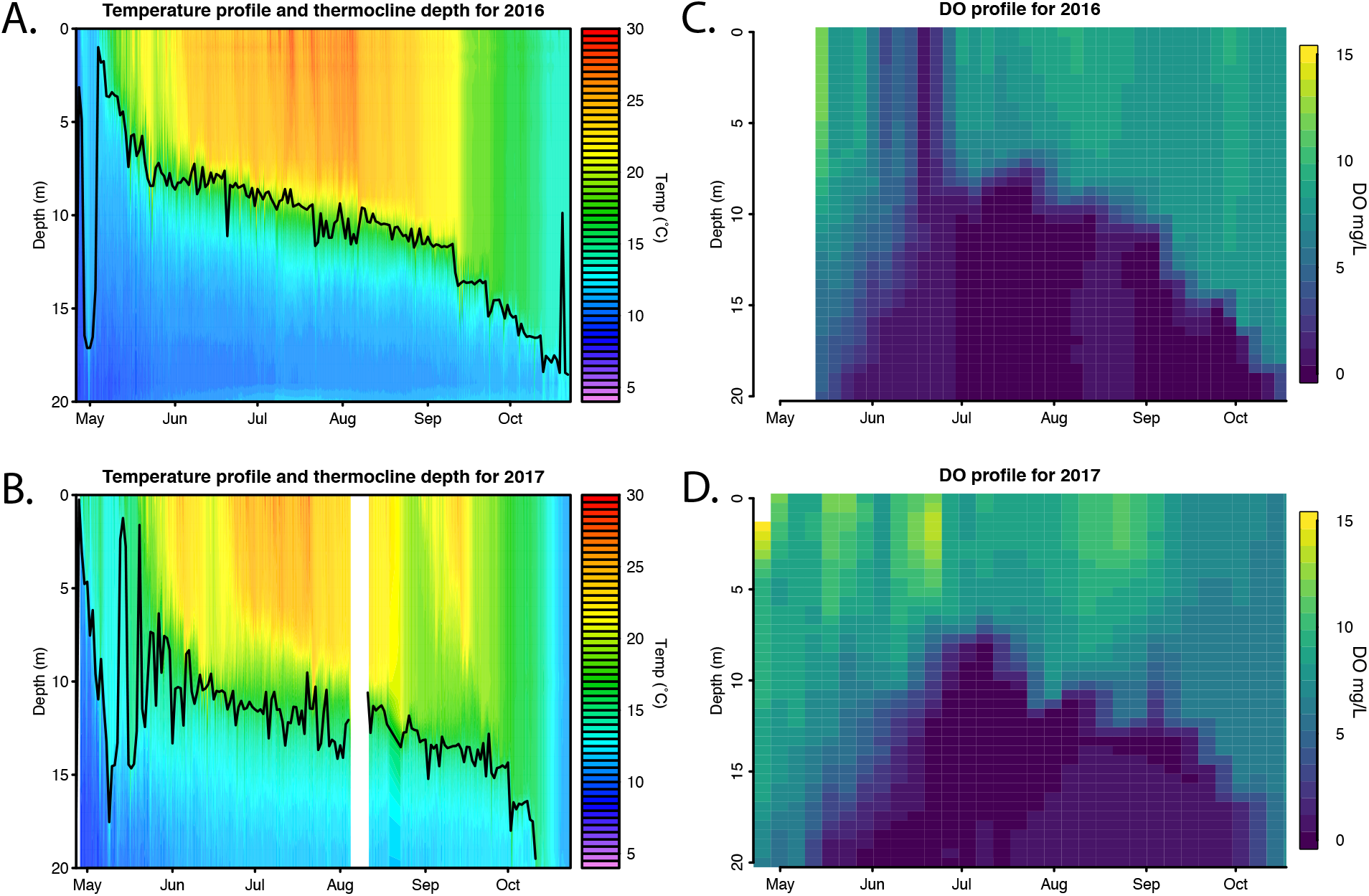
Temperature and Dissolved Oxygen profile of 2016 and 2017. These plots were compiled from data gathered from a buoy that is positioned spring through fall at the sampling site. In the temperature profiles A and B, The line represents the thermocline position at a given point in time. The thermocline gradually moves down as the year progresses preceding the fall mixing event. The thermocline is in similar positions in 2016 and 2017. Graphs C and D are the corresponding DO profiles. DO profiles are very similar between the years.

**Figure S3.**
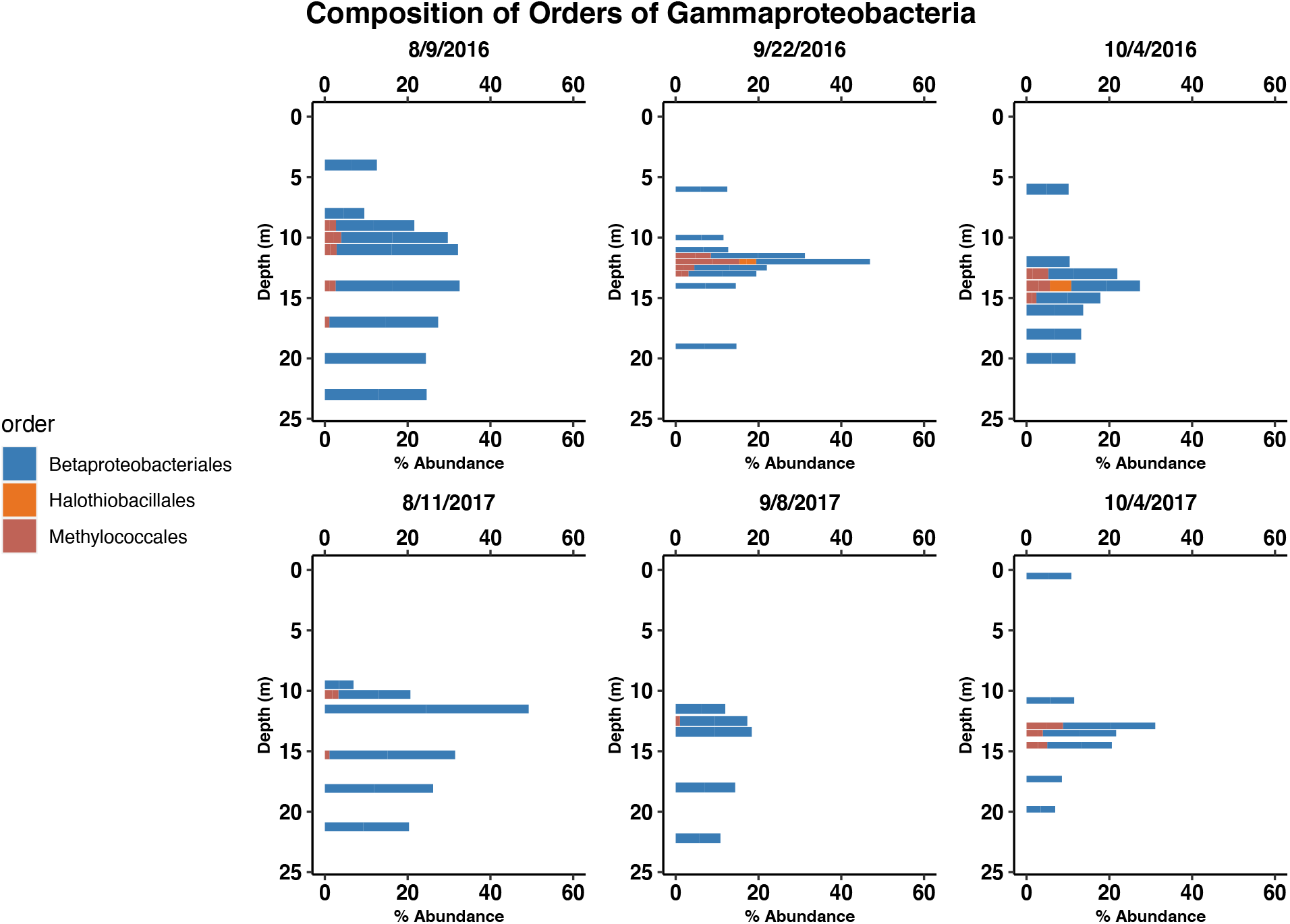
Composition of Class Gammaproteobacteria. Bar charts displaying the distribution and the composition of orders in class Deltaproteobacteria for each sample. Samples for each date were taking at several discreet depths above, at and below the oxycline. Order Betaproteobacteriales dominate the class at the epilimnion and hypolimnion.

**Figure S4.**
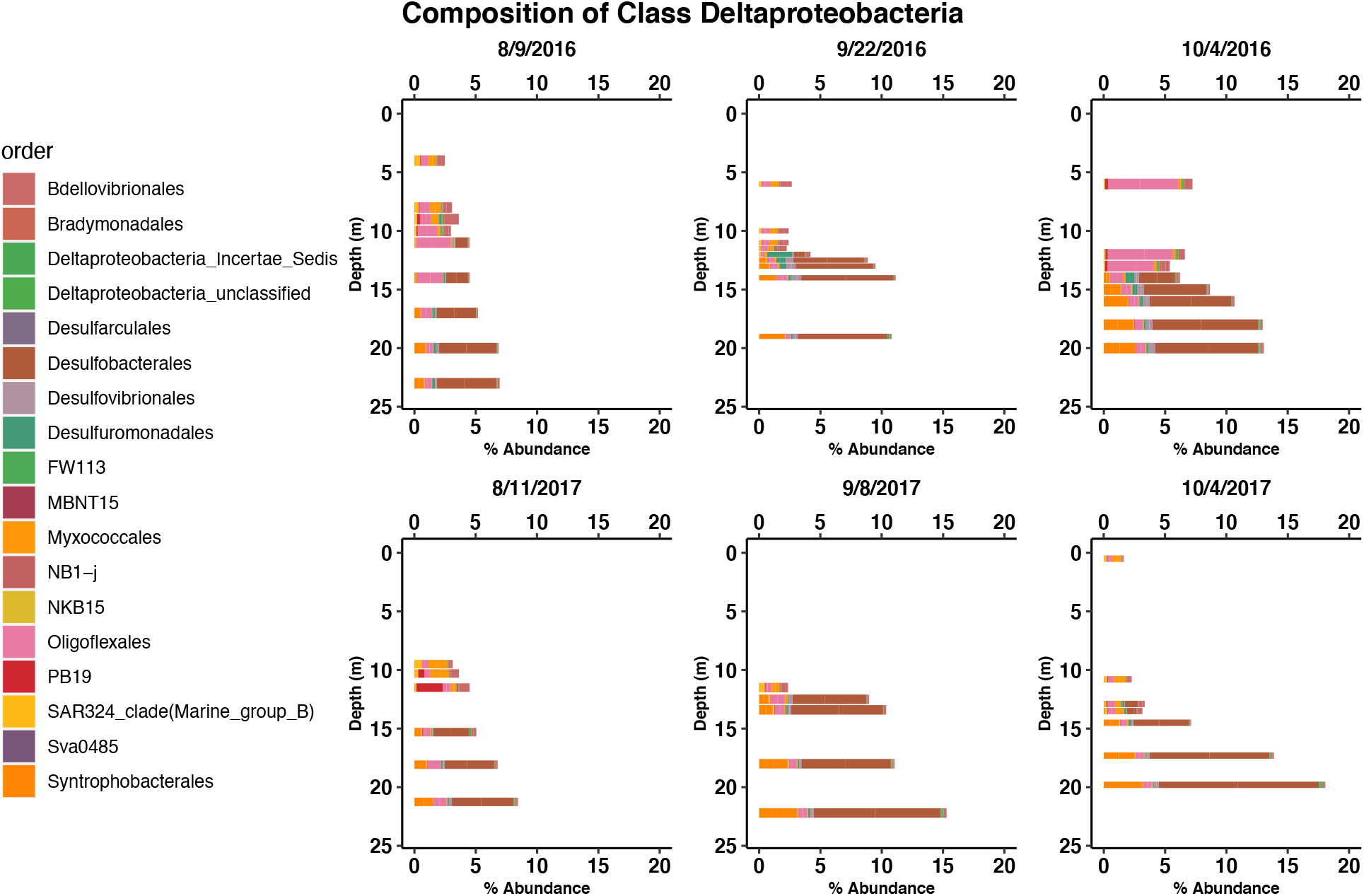
Composition of class Deltaproteobacteria. Bar charts displaying the distribution and the composition of orders in class Deltaproteobacteria for each sample. Samples for each date were taken at several discreet depths above, at and below the oxycline. Desulfobacterales are the most prevalent in the hypolimnion correlating with the increase of sulfide.

**Figure S5.**
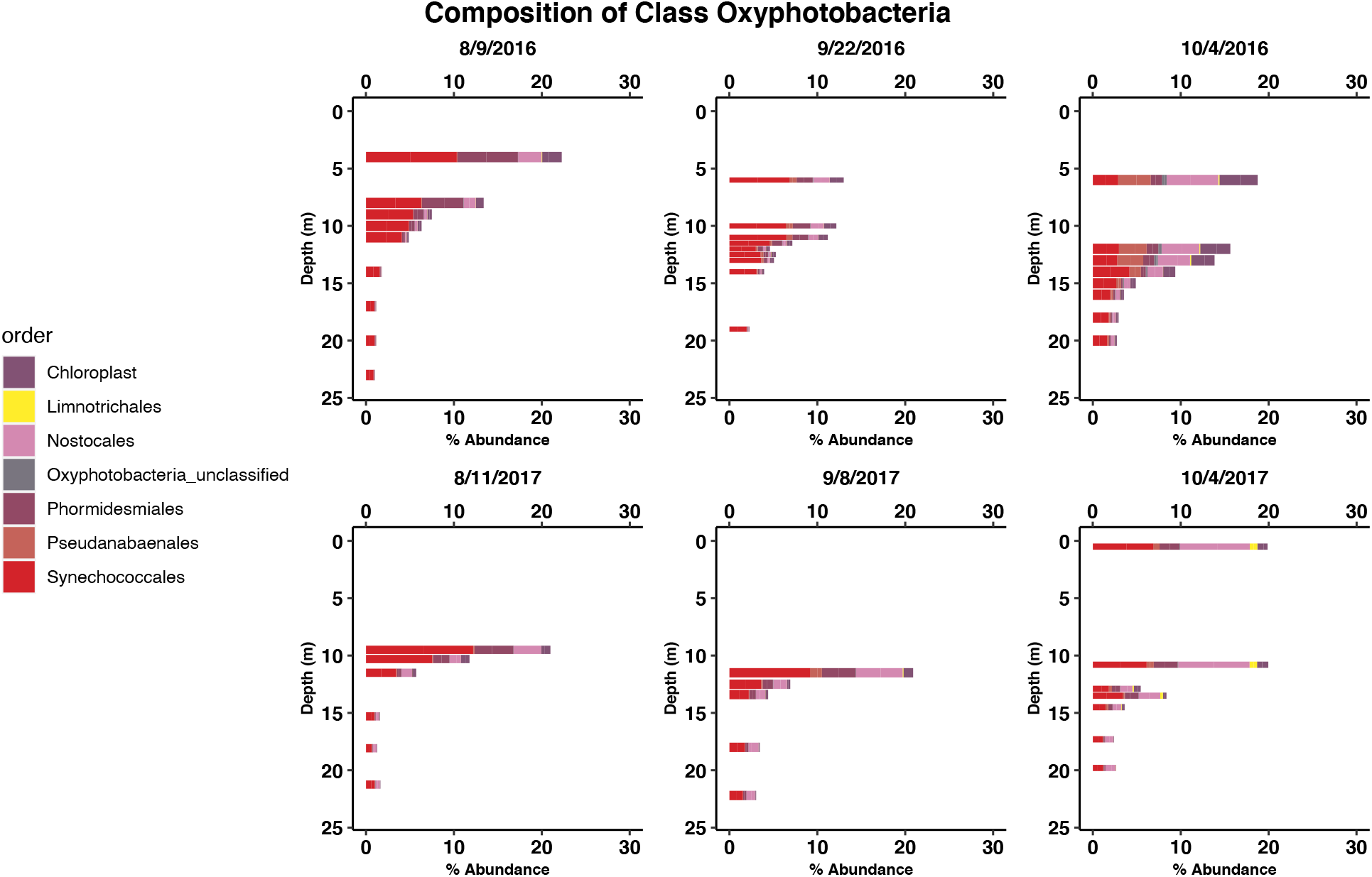
Composition of class Oxyphotobacteria. Bar charts displaying the distribution and the composition of orders in class Oxyphotobacteria for each sample. Samples for each date were taken at several discreet depths above, at and below the oxycline. Although Oxyphotobacteria are primarily in the epilimnion, surprisingly some Synechococcales are capable of survival at the depths of lake Mendota.

**Figure S6.**
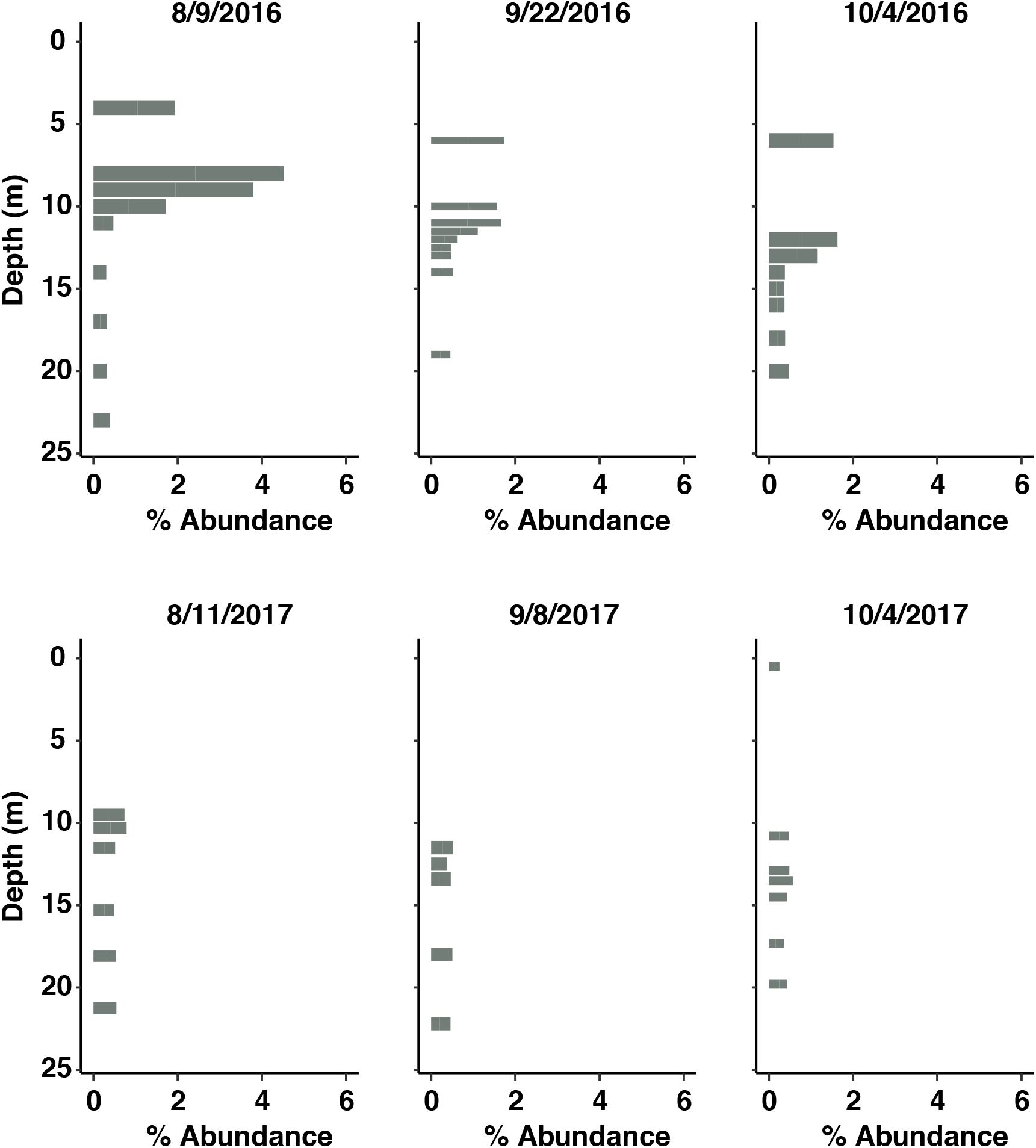
Vertical Distribution of Ignavibacteria. Bar charts displaying the distribution and relative abundance of class Ignavibacteria. Samples for each date were taken at several discreet depths above, at and below the oxycline. This graph shows the relative abundance of class Ignavibacteria with respect to all ASVs. Ignavibacteria tend to spike in the lower epilimnion corresponding with an uptick in manganese and waning DO.

**Figure S7.**
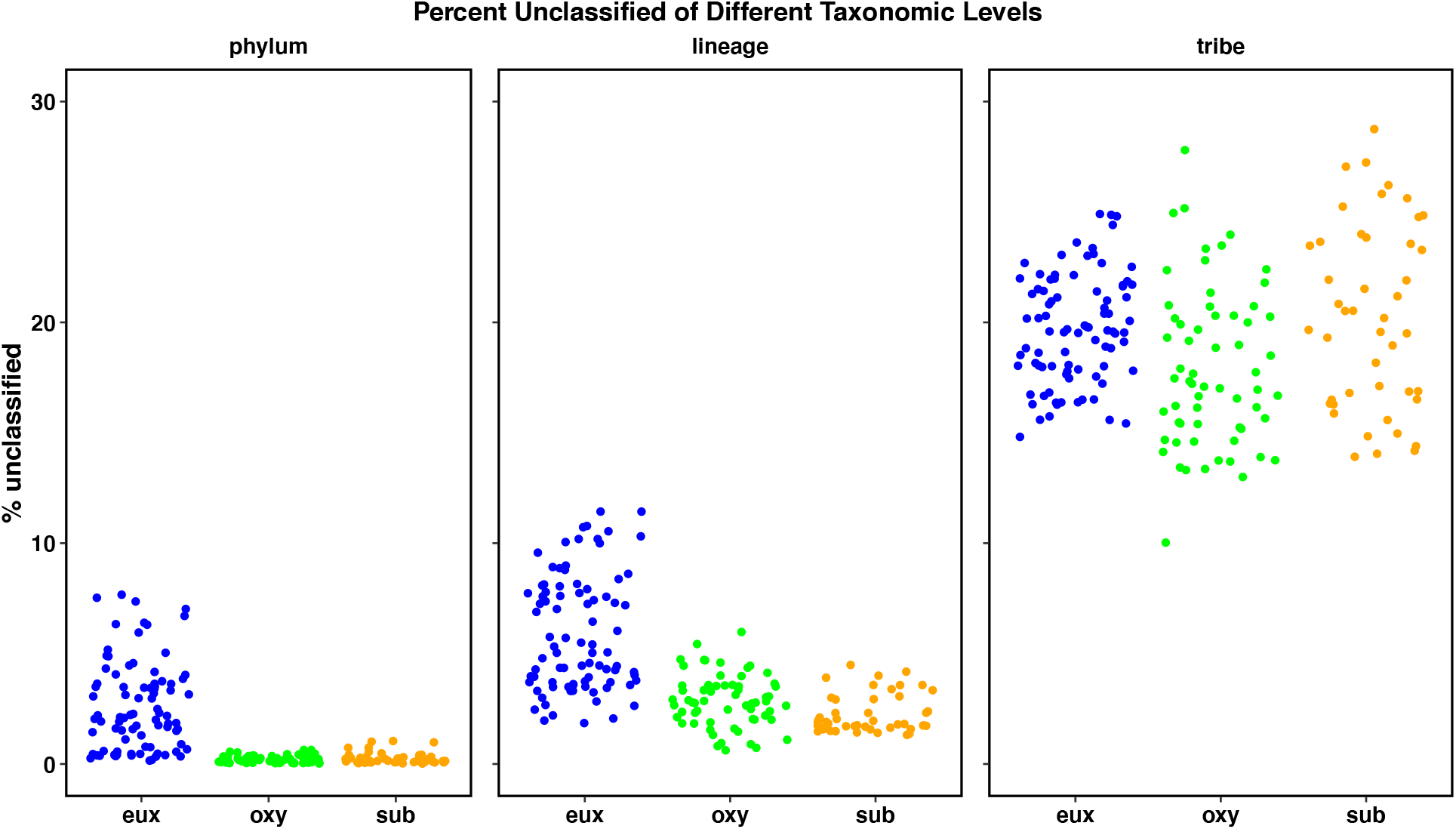
Percent of Unclassified Sequences in each Sample Strip plots showing the percent of unclassified sequences in each sample for three phylogenetic levels.

**Figure S8.**
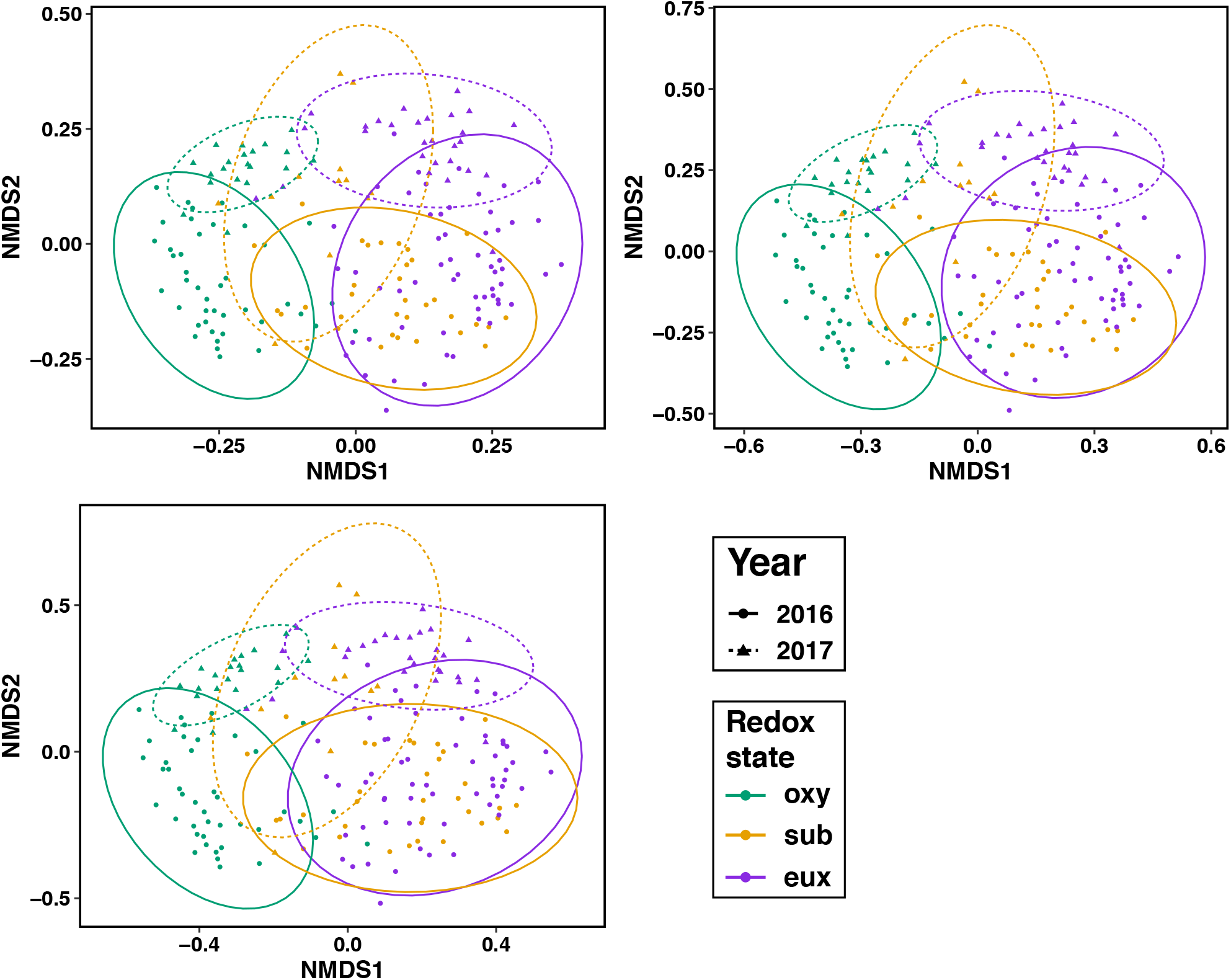
NMDS ordinations at different ASV cutoffs. Nonmetric Multidimensional Scaling (NMDS) using Bray-Curtis distance to display dissimilarity between each sample’s ASVs. Ellipses represent the clustering of samples by redox state (color) and year (line type) using a 95% confidence level. The greater than symbol along with the number at the top of each ordination is referring to the cutoff of that ordination. For example >100 means only ASVs that appear more that 100 times were used in the ordination. All three graphs look very similar with only the 2016 sub ellipse changing position a little bit.

## Notes

https://github.com/robertmarick/Mendota20162017

https://osf.io/p92gn/

### Competing Interest Statement

The authors have declared no competing interest.

